# A temperature sensitive mutation in the CstF77 subunit of the polyadenylation complex reveals the critical function of mRNA 3’ end formation for a robust heat stress response in plants

**DOI:** 10.1101/2021.10.31.466691

**Authors:** Minsoo Kim, John Swenson, Fionn McLoughlin, Elizabeth Vierling

## Abstract

**Background:** Heat Shock Protein 101 (HSP101) in plants and orthologs in bacteria (Caseinolytic peptidase B, ClpB) and yeast (Hsp104) are essential for thermotolerance. To investigate molecular mechanisms of thermotolerance involving HSP101, we performed a suppressor screen in *Arabidopsis thaliana* of a semi-dominant, missense HSP101 allele, *hot1-4* (A499T). Plants carrying the *hot1-4* mutation are more heat-sensitive than an HSP101 null mutant (*hot1-3*), indicating the toxicity of *hot1-4* allele.

**Results:** We report that one suppressor (*shot2, suppressor of hot1-4 2*) has a temperature-sensitive, missense mutation (E170K) in the CstF77 (Cleavage stimulation factor 77) subunit of the polyadenylation complex, which is critical for 3’ end maturation of pre-mRNA. RNA-Seq analysis of total RNA depleted of ribosomes reveals that heat treatment causes transcriptional readthrough events in *shot2*, specifically in highly heat-induced genes, including the toxic *hot1-4* gene. In addition, failure of correct transcript processing leads to reduced accumulation of many HSP RNAs and proteins, suppressing heat sensitivity of the *hot1-4* mutant, due to reduction of the toxic mutant HSP101 protein. Notably, the *shot2* mutation makes plants more sensitive to heat stress in the HSP101 null (*hot1-3*) and wild-type backgrounds correlated with the reduced expression of other heat-inducible genes required for thermotolerance.

**Conclusions:** Our study reveals the critical function of CstF77 for 3’ end formation of mRNA during heat stress, as well as the dominant role of HSP101 in dictating the outcome of severe heat stress in plants.

## Background

Exposure of organisms to elevated temperatures leads to dramatic reprogramming of gene expression, including transcriptional upregulation of heat shock proteins (HSPs) and other proteins necessary to protect living organisms from heat damage (1). HSPs act as molecular chaperones to protect and/or rescue critical client proteins from denaturation and aggregation (2). HSP synthesis is one of the most important steps required for organisms, including plants, to acquire thermotolerance during heat acclimation processes (3), and HSP synthesis depends on the many steps involved in generating correctly processed, mature mRNAs.

In eukaryotes, production of mRNAs that can engage in translation requires multiple steps starting with transcription into precursor messenger RNAs (pre-mRNAs) in the nucleus by RNA polymerase II. High temperatures are known to trigger activation of transcription factors including HSFs that bind to the promoter regions of heat stress responsive genes including HSPs (1,4,5). RNA polymerase II then initiates transcription of the target genes. Pre-mRNAs must then be processed into mature mRNAs by multiple steps including 5’ capping, splicing and polyadenylation of the 3’ end before being transported to the cytoplasm for translation. Polyadenylation of the 3’ end is achieved by cleavage of pre-mRNAs at a specific location downstream of a polyadenylation signal and the addition of a poly(A) tail at the cleavage site (6,7). Studies in yeast and mammals have identified many protein factors required for 3’ end processing, including the cleavage and polyadenylation specificity factor (CPSF), cleavage stimulation factor (CstF), and poly(A) polymerase (PAP) (6,8). These core factors involved in mRNA 3’ end formation are highly conserved in plants (9,10). Understanding how these core proteins function in plants is important because mRNA 3’ end processing can affect gene expression at the levels of mRNA stability, nuclear export, transcript localization, and translation efficiency as studied in other organisms (11–14). Defining the impact of mRNA 3’ end processing on transcription termination and mRNA stability during heat or other stress conditions is also relevant to understanding critical mechanisms of stress tolerance.

Among the HSP genes induced by high temperature in plants, HSP101 has been clearly documented by genetic analysis to be required for heat acclimation leading to survival of severe heat stress. HSP101 belongs to the Hsp100/ClpB chaperone family of hexameric AAA+ (ATPases Associated with various cellular Activities) proteins, which play essential roles in acquired thermotolerance not only in plants, but also in bacteria and yeasts (15–18). These chaperones act by dissociating protein aggregates and helping denatured proteins refold into native structures using ATP as energy (19,20). The importance of this protein in plants was first demonstrated by the identification of loss-of-function mutations in *Arabidopsis thaliana* HSP101 (*hot1* mutants), which make plants particularly sensitive to extreme heat stress (17). *A. thaliana* HSP101, like other Hsp100/ClpB protein disaggregases, is composed of an N-terminal domain, two conserved AAA+ nucleotide-binding domains (NBD1 and NBD2) and a characteristic coiled-coil middle domain (MD) inserted near the C-terminal end of NBD1 (21). Hsp70 regulates activities of Hsp100/ClpB by interacting with the MD (22–24). Mutations in the MD of the orthologous yeast Hsp104 and *Escherichia coli* ClpB exert diverse effects on the protein’s function suggesting the MD has crucial roles in regulating the activity of this protein family (22,25–27). Notably, some MD mutations cause constitutive derepression of protein activity leading to toxicity *in vivo* (25,27,28). A gain-of-function mutation in the MD of *A. thaliana* HSP101 (*hot1-4*; A499T) results in plants that cannot survive acclimation temperatures that induce HSP101 - temperatures that are not otherwise lethal to HSP101 null plants (*hot1-3* mutants) and that are necessary to acclimate wild-type plants to survive severe heat stress (29). Sensitivity of *hot1-4* plants to heat stress acclimation temperatures indicates that the *hot1-4* mutant protein is toxic, as is seen for the MD mutations in other organisms.

To understand further the function of HSP101 in mechanisms of plant thermotolerance, we performed suppressor screening of ethyl methanesulfonate-mutagenized seeds of *A. thaliana* carrying the *hot1-4* (A499T) missense mutation in HSP101. In addition to multiple intragenic suppressors (Lee et al., 2005), we also identified extragenic suppressors. We previously reported identification of the extragenic suppressor *shot1* (*suppressor of hot1-4 1*), which is a bypass suppressor mutant with defects in mitochondrial function (30,31). Here, we report the isolation of three additional extragenic suppressor mutants (*shot2, 3* and *4*) and the identification of the *shot2* mutation in a gene encoding the CstF77 protein (AT1G17760), a component of the polyadenylation complex involved in formation of the 3’ end of mRNA. We demonstrate that the *shot2* mutation (E170K) is a temperature-sensitive allele of CstF77 that loses activity at elevated temperatures. The *shot2* mutant allele gave us the opportunity to investigate the effects of a defective polyadenylation complex on gene expression and thermotolerance. RNA-Seq data reveal that *shot2* negatively affects transcription termination of highly heat-inducible genes as evidenced by detection of transcript readthrough leading to reduced transcript levels of genes critical to thermotolerance. Further, in the *hot1-4* mutant background, the *shot2* mutation causes reduction of the toxic mutant HSP101 (A499T) protein, which enables *hot1-4* plants to better tolerate acclimation heat stress temperatures that induce HSP101. However, in the HSP101 null *hot1-3* mutant or wild-type backgrounds, because the *shot2* mutation also causes reduction of many heat-inducible transcripts critical for thermotolerance, the plants more sensitive to heat stress. Our study highlights the conserved function of CstF77 in RNA processing and emphasizes the central role of HSP101 in the protein quality control network activated upon heat stress.

## Results

### Extragenic suppressors of *hot1-4*

The degree of heat sensitivity of dark grown seedlings can be quantitatively measured by hypocotyl growth inhibition after heat stress, which provides an assay to identify mutants with altered heat stress sensitivity (17). When *A. thaliana* seedlings are treated at 38 °C for 1.5 h followed by two or more hours at optimal growth temperatures, the plants accumulate HSPs and can become acclimated and survive treatments at a more severe, 45 °C temperature (17,29). However, because 38 °C treatment of seedlings carrying the HSP101 *hot1-4* allele produces a toxic, mutant HSP101 protein, *hot1-4* seedlings do not grow after a 38 °C treatment, which is an otherwise permissive treatment for wild type and HSP101 null plants(29). We therefore screened for suppressors of the *hot1-4* mutant allele by subjecting three-day-old, dark grown M2 seedlings of ethyl methanesulfonate-mutagenized *hot1-4* to a heat treatment at 38 °C for 2 h and rescued surviving seedlings. We report the identification of three extragenic suppressors (*“shot”* mutants: **s**uppressor of ***hot**1*) of *hot1-4* and present detailed analysis of one suppressor, *shot2*.

The three *shot* mutants show the ability to suppress the toxic effect of the *hot1-4* allele after a 38 °C treatment for 3 h (Fig. 1a), a condition that is permissive for growth of the wild-type Columbia-0 (Col-0) and for the HSP101 null mutant, *hot1-3*, as shown previously (29). The level of HSP101 accumulation was examined, since reduced levels of the dominant negative, toxic *hot1-4* mutant protein could result in the suppressor phenotype. As shown in Fig. 1b, HSP101 is reduced in *hot1-4 shot2* compared to the wild type and *hot1-4*, while the other two suppressors have HSP101 levels similar to wild type. Therefore, the suppressor phenotype of *hot1-4 shot2* may result from the decreased level of the toxic HSP101 hot1-4 protein.

**Fig. 1.**
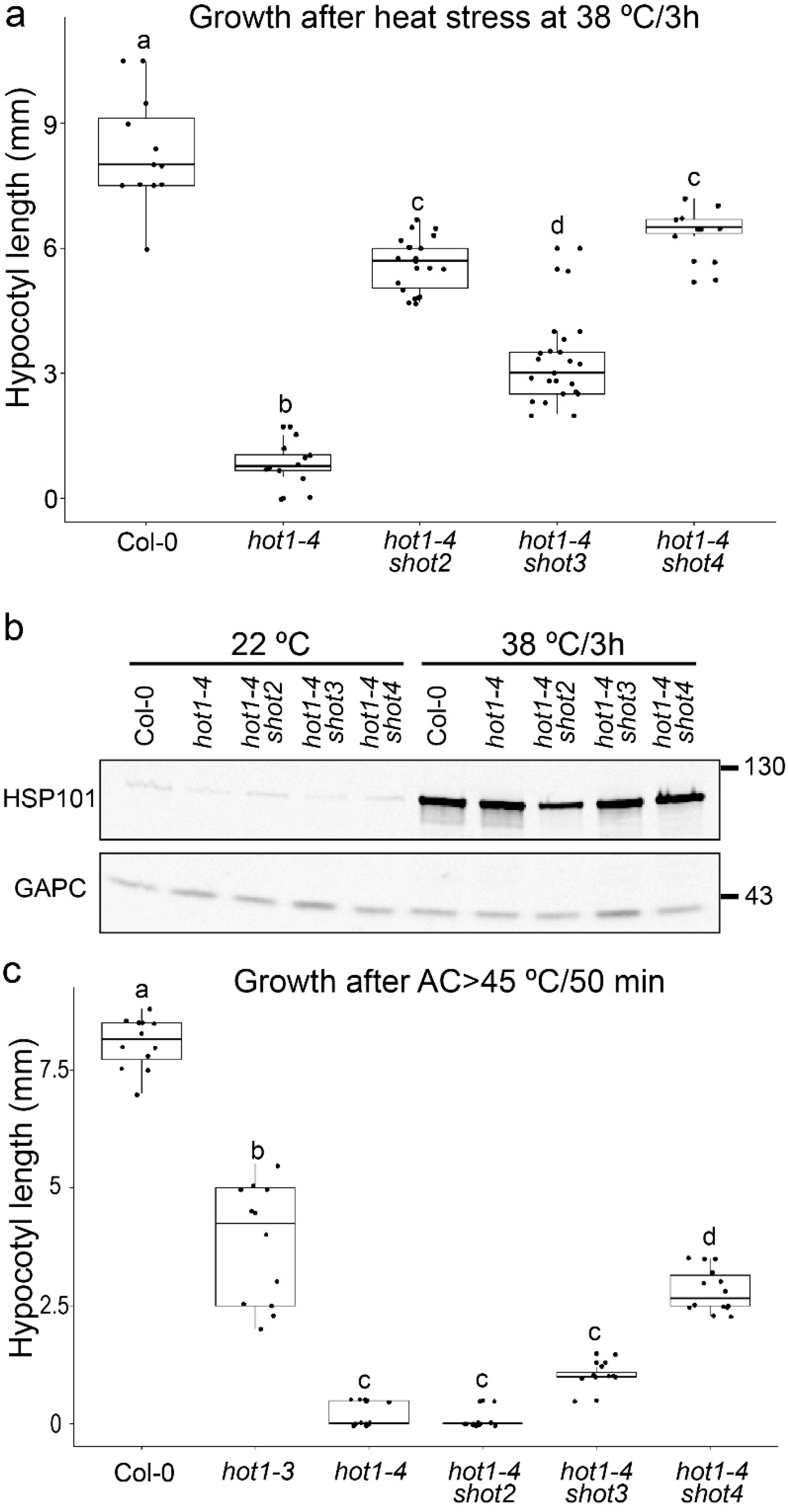
Heat stress phenotype and immunoblot analysis of extragenic suppressors of *hot1-4*. (a) Three suppressor mutants were isolated. The degree of suppression was measured by hypocotyl elongation of dark-grown seedlings after heat treatment at 38 °C for 3h. Different letters indicate significant differences (p<0.01) by one-way ANOVA followed by Tukey’s post hoc test. (b) HSP101 protein levels were checked by immunoblot analysis of samples from three-day-old dark-grown seedlings with or without heat treatment at 38 °C for 3h. Immunoblot against anti-GAPC (glyceraldehyde-3-phosphate dehydrogenase, cytosolic) is shown as a loading control. (c) Heat stress tolerance was measured by hypocotyl elongation after 45 °C for 50 min following acclimation treatment (AC = 38 °C for 1.5h, 22 °C for 2h). Different letters indicate significant differences (p<0.01) by one-way ANOVA followed by Tukey’s post hoc test. The hypocotyl elongation assays in (a) and (b) were repeated more than three times with similar results.

The suppressor mutants were then tested in an acquired thermotolerance assay (heat acclimation (AC) at 38 °C/1.5 h plus 22 °C/2 h, followed by heat stress at 45 °C/50 min), where the 38 °C treatment leads to ability to survive the subsequent 45 °C and for which the function of HSP101 is required (17). In this assay all suppressor mutants were more heat-sensitive than the HSP101 null *hot1-3* mutant (Fig. 1c), which still exhibits some hypocotyl elongation. In total, these heat sensitivity data indicate that none of the suppressor mutants has restored wild-type HSP101 function to hot1-4, but rather have in some way reduced the negative effect of hot1-4.

We chose to examine the *shot2* suppressor in more detail because of the strong suppression phenotype at 38 °C and an interest in understanding how the mutation controlled HSP101 levels. To determine the genetic inheritance of the *shot2* mutation, *hot1-4 shot2* was back-crossed to the parental *hot1-4* mutant. The F1 plants were all heat-sensitive at 38 °C like *hot1-4* suggesting that the mutation is recessive (Additional file 1: Fig. S1a). A quarter of the F2 plants showed the suppressor phenotype (Additional file 1: Fig. S1b) indicating that the phenotype is caused by a recessive mutation at a single genetic locus.

### The *SHOT2* gene encodes CstF77, a component of the pre-mRNA 3 end processing complex

To identify the causal mutation of the *shot2* suppressor phenotype, an F2 mapping population was created by crossing *hot1-4 shot2* (Columbia-0 background; Col-0) with the Landsberg *erecta* (L*er*) ecotype harboring the *hot1-4* mutation. The *hot1-4* mutation had been introgressed into L*er* by backcrossing nine times to Col-0 *hot1-4*. From the mapping population, we selected 100 individual F2 plants with the suppressor phenotype and selfed them to obtain the F3. The suppressor phenotype was confirmed by assaying F3 seedlings from each of the 100 F2 lines, and then eight F3 seedlings from each F2 line were grown and pooled for whole genome sequencing. We located the region of interest to the upper arm of chromosome 1 (Fig. 2a). By examining the frequency of L*er*-specific single nucleotide polymorphisms (SNPs) in the F3 genome sequencing data, we further narrowed the region to four candidate genes (Additional file 2: Table S1). Based on the predicted functions and expression patterns of the candidates, we chose the *CstF77* gene (AT1G17760) for complementation tests.

**Fig. 2.**
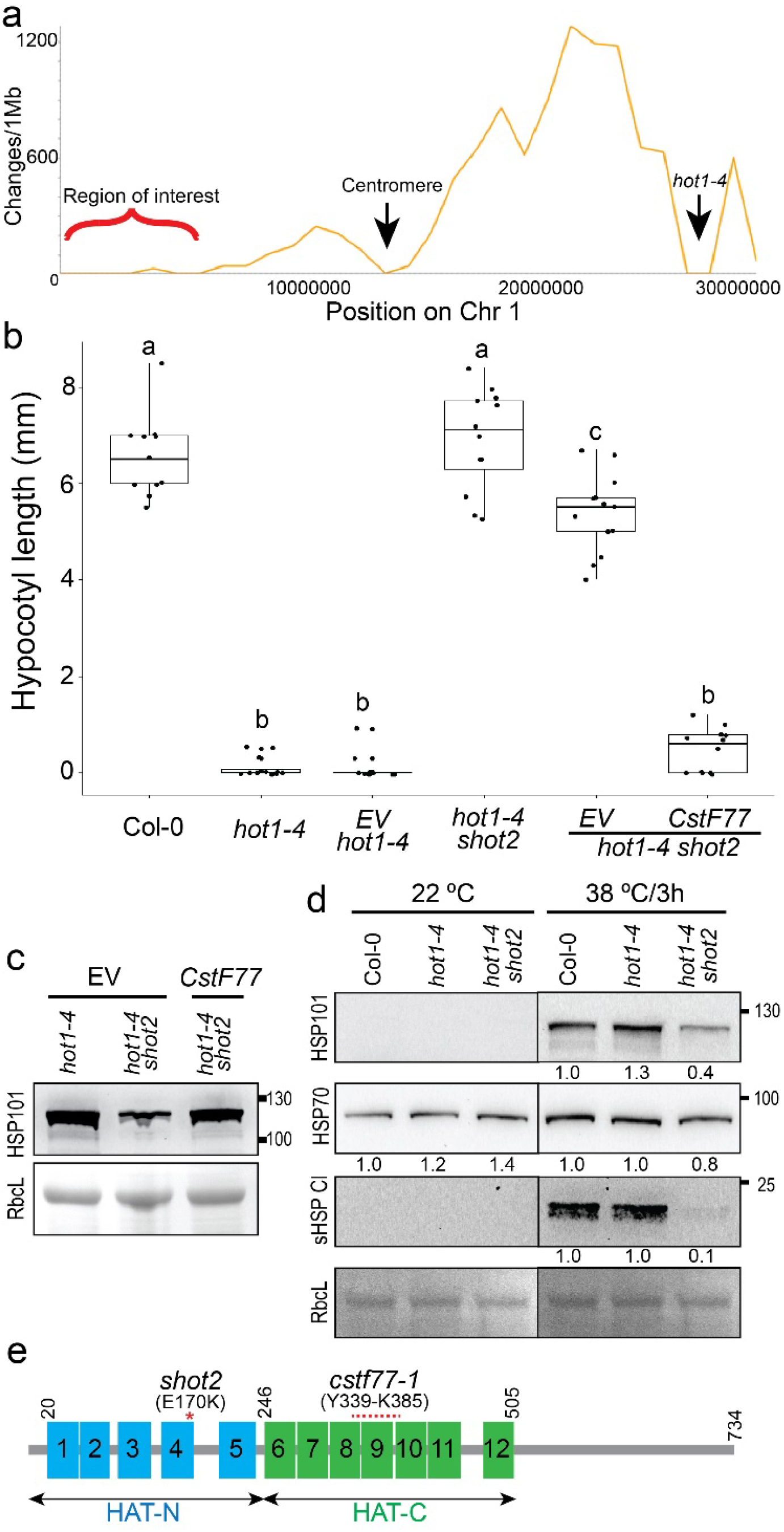
A mutation in the *CstF77* gene is responsible for the suppressor phenotype of the *shot2* mutant. (a) DNA sequence changes in the F3 mapping population were plotted along chromosome 1. The causal mutation for the suppressor phenotype is located on the upper arm of chromosome 1 as indicated. (b) T1 transgenic plants carrying the wild-type *CstF77* gene showed recovery of the heatsensitive phenotype in *hot1-4 shot2*. Hypocotyl length of dark-grown seedlings was measured after heat treatment at 38 °C for 3h. Different letters indicate significant differences (p<0.01) by one-way ANOVA followed by Tukey’s post hoc test. The assay was performed twice with similar results. EV: Empty vector control. (c) Immunoblot shows that the introduction of the wild-type *CstF77* gene recovers accumulation of HSP101. Protein was extracted from leaves of 3-week-old T1 plants of the indicated genotypes. Ponceau-S stained rubisco large subunit (RbcL) is shown as a loading control. (d) Immunoblot analysis of HSP101, HSP70 and cytosolic class I sHSPs (sHSP CI). The *hot1-4 shot2* mutant accumulates less HSP101 and cytosolic class I sHSPs. Ponceau-S stained rubisco large subunit (RbcL) is shown as a loading control. (e) Domain organization of the SHOT2/CstF77 protein. The HAT (Half-A-TPR)-N and −C domains contain the first five HAT motifs (blue) and the other seven HAT motifs (green), respectively. Position of the *shot2* mutation and the *cstf77-1* allele discussed in the text are denoted in red.

A 9009 bp genomic DNA construct encompassing the promoter (2032 bp upstream of the start codon) and terminator (913 bp downstream of the stop codon) regions of the *CstF77* gene was introduced into *hot1-4 shot2* plants. T1 transgenic plants carrying the *CstF77* gene became sensitive to heat stress like *hot1-4*, while control T1 plants containing the empty vector were still resistant like *hot1-4 shot2*, confirming that the *shot2* mutation in the *CstF77* gene is responsible for the suppressor phenotype (Fig. 2b). Immunoblot analysis confirmed the restoration of HSP101 protein levels in the transgenic plants carrying *CstF77* (Fig. 2c). The *shot2* mutation also negatively affects the accumulation of cytosolic class I small HSPs (sHSPs) indicating that the function of CstF77 may be required for accumulation of other HSPs (Fig. 2d). The level of cytosolic HSP70s was also reduced after heat treatment in *shot2* mutant plants, although the extent of the decrease is difficult to detect because the antibody reacts with all five cytosolic HSP70s including constitutively expressed isoforms (Fig. 2d).

In eukaryotes, 3’ end processing of messenger RNA precursors (pre-mRNAs) requires a large multi-subunit protein complex, which includes the cleavage and polyadenylation specificity factor (CPSF), cleavage stimulation factor (CstF), and poly(A) polymerase (PAP) (6,8). Pre-mRNAs are cleaved at a specific location at the 3’ end by CPSF with the help of CstF, and a poly(A) tail is added by PAP before the mRNAs are exported to the cytoplasm for translation. The CstF complex is composed of CstF77, CstF64 and CstF50, and it helps stabilize CPSF to allow cleavage of pre-mRNAs (32,33). CstF77 is evolutionary conserved in eukaryotes from yeast to humans and plants (6,34,35). CstF77 has multiple interactions with other polyadenylation factors including the other CstF subunits, as well as CPSF160 and symplekin (36,37). The N-terminal, approximately two thirds of CstF77 comprises tandem arrays of 12 HAT (Half-a-TPR) motifs, which are divided into HAT-N (HAT1-5 motifs) and HAT-C (HAT6-12 motifs) domains, while the final third of the protein is mostly disordered, highly flexible and not visible in available high resolution structures (Fig. 2e) (35,38–40). The HAT-C domain is important for homodimerization (38–40) and interaction with CPSF160 (40), while interaction with CstF64 and CstF50 appears mediated by the flexible C-terminal region (41,42). The *shot2* mutant carries a missense mutation in the HAT-N domain at an evolutionary conserved residue in the fourth HAT motif (E170K) (Fig. 2e and Additional file 1: Fig. S2). The corresponding residue in human CstF77, E194, appears hydrogen-bonded with R169 (40), which is also conserved in *A. thaliana* (R145) (Additional file 1: Fig. S3). Disruption of this H-bond by the *shot2* mutation may be the basis for the temperature sensitivity of the mutant protein. The first mutant allele of CstF77 identified in *A. thaliana, cstf77-1*, carries a deletion from Y339 to K385 in the HAT-C domain, and was reported as one of the suppressors of overexpressed *FLOWERING CONTROL LOCUS A* (FCA) (34). A T-DNA insertional knockout allele, *cstf77-2*, was initially reported to be lethal in the same study (31), but later found to be viable, although it has serious growth defects (43). Importantly, the *shot2* mutation has no negative effect on plant growth under standard growth conditions, indicating that the *shot2* mutation does not cause any obvious defects to CstF77 function in the absence of temperature stress (Additional file 1: Fig. S4). These data indicate that the *shot2* mutation is a temperature-sensitive allele, which loses its activity only at high temperatures.

### Heat stress treatment of *shot2* leads to readthrough transcription and reduced expression of heat-inducible genes

The temperature-sensitive *shot2* mutation in the *CstF77* gene is expected to cause defects in 3’ end processing not only of *HSP101* mRNA, but also of many other mRNAs including *sHSPs*, which are reduced at the protein level (Fig. 2d). We also predicted that the *shot2* mutation would cause lower transcript levels due to polyadenylation defects. Therefore, we performed RT-PCR analysis of heat-inducible genes after heat-treatment at 38 °C for 3h. As expected, transcript levels of *HSP101, HSP17.6A* and *UBQ10* were significantly reduced in the *hot1-4 shot2* mutant (Additional file 1: Fig. S5).

To obtain a transcriptome-wide picture of defects caused by the *shot2* mutation, we performed RNA-Seq analysis comparing *hot1-4 shot2* to *hot1-4* plansts either with or without a heat treatment (38 °C/3h) in three biological replicates each. Since the *shot2* mutation is expected to cause alterations in polyadenylation, ribosomal RNA depletion rather than poly-A enrichment was performed before constructing directional libraries. About 40 million 75 bp single-end reads were mapped to the *A. thaliana* genome (TAIR10) using HISAT2 (44). About 70 % of reads mapped to multiple locations, mostly to rRNAs, which is not surprising because of rRNA abundance even after rRNA depletion procedures. Nevertheless 10-15 million reads from each sample were uniquely mapped to the genome and used for further analyses.

A multi-dimensional scaling (MDS) plot of the RNA-Seq expression profiles shows that there are virtually no differences in transcripts between *hot1-4* and the suppressed *hot1-4 shot2* genotype under normal growth conditions (Fig. 3a), confirming our prediction that *shot2* is a temperature-sensitive mutation and reflecting the absence of phenotypic differences in the *hot1-4 shot2* mutant compared to wild type under optimal growth conditions. Using significance cutoff values of FDR < 0.01 and |log2FC| >1, only 26 out of a total of 17320 genes were identified as differentially expressed between the two genotypes under normal growth conditions (Additional file 2: Table S2). There was no significant enrichment of GO terms from this set of differentially expressed genes. Heat stress resulted in dramatic changes in the transcriptome of both genotypes; 30.7% and 33.7% of detected transcripts in *hot1-4* and *hot1-4 shot2*, respectively, were differentially expressed by the stress treatment. For *hot1-4*, this comprised increased transcripts of 2785 genes and decreased transcripts of 2541 genes (Additional file 2: Table S3), while the suppressed *hot1-4 shot2* mutant showed a total of 2798 genes increased, and 3071 genes decreased (Additional file 2: Table S4). However, as evidenced by the major separation of the two genotypes for the heat stressed samples on the MDS plot (Fig. 3a), when the RNA-Seq profiles after heat treatment of the two genotypes were compared, more than twice as many transcripts were decreased (1356) rather than increased (584) by the presence of the *shot2* mutation compared to plants carrying the *hot1-4* mutation alone (Additional file 2: Table S5). Therefore, plants carrying the *shot2* mutation are capable of a major transcriptional response to heat stress, but the mutation significantly alters the transcriptome profile.

**Fig. 3.**
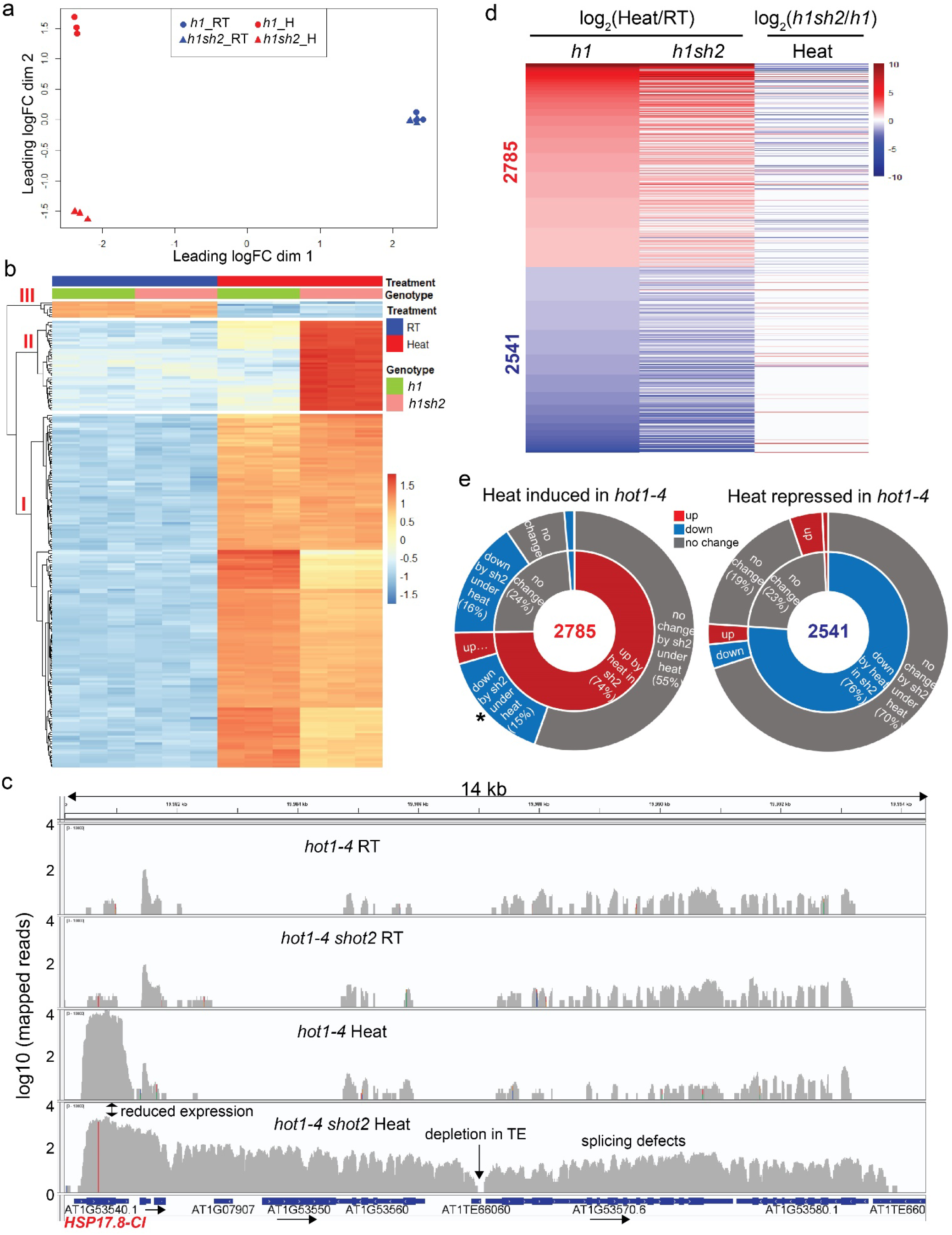
Most heat-inducible genes accumulate lower transcripts in *hot1-4 shot2* compared to *hot1-4*. (a) Multi-dimensional scaling plot of RNA-Seq expression profiles in two dimensions. The *hot1-4* (*h1*) and *hot1-4 shot2* (*h1sh2*) were treated with (H) or without (RT) heat stress (38 °C/3h). There are three biological replicates for each group. (b) Heat map of the top 200 most variable genes from the RNA-Seq experiment. The genes were grouped into three clusters (I-III) as indicated in the cladogram. Expression is scaled to z-scores of log2 (counts per million). (c) Mapped reads on HSP17.8-CI and its downstream region showing abundant readthrough reads in *hot1-4 shot2* plants under heat treatment (log10 scale). The HSP17.8-CI gene is oriented from left to right. Genes with the same orientation are shown with arrows. TE: transposable element. The vertical red line indicates a missense mutation (A65V) in *hot1-4 shot2* introduced by mutagenesis. (d) The 5326 genes differentially expressed in *hot1-4* in response to heat treatment (FDR < 0.01, |log2FC| >1, left column) were tracked in *hot1-4 shot2* samples upon heat treatment (compared to *hot1-4 shot2-RT*, middle column) and under heat treatment (compared to *hot1-4*-Heat, right column). (e) Pie charts showing proportions of differentially expressed genes as shown in (d). 5326 genes expressed differentially in *hot1-4* by heat were separated into two pies (2785 genes upregulated on the left and 2541 genes downregulated on the right). The inner and outer pies represent proportions in the middle column and right column in (d), respectively.

To investigate further transcriptome differences between the two genotypes with or without the heat treatment, the top 200 genes with most variation were chosen to generate the heat map in Fig. 3b. The heat map defines three major gene clusters (See gene list in Additional file 2: Table S6). The first cluster contains the most genes (154), which were upregulated by heat in both genotypes. This cluster contains many HSP genes including HSP101, HSP70s, and sHSPs. These genes are heat-induced in *hot1-4 shot2*, but show readthrough reads downstream of their typical transcription termination sites (TTS) as illustrated for AT1G53540 (HSP17.8) in Fig. 3c. Also, the numbers of read counts in the genic region (i.e., expression values) are significantly reduced in *hot1-4 shot2*, which is consistent with the RT-PCR experiments (Additional file 1: Fig. S5). The second cluster, with 39 genes, contains genes up-regulated by heat specifically in *hot1-4 shot2*, of which 38 are located downstream of heat-induced genes. Nineteen of these upstream genes are canonical HSPs that affect 26 of the 39 up-regulated downstream genes. Four different HSP70 genes and 10 different sHSPs show downstream reads as illustrated for the sHSP *HSP17.8* (AT1G53540) (Fig. 3c). Their increased transcript levels reflect apparent transcription termination defects of upstream genes caused by the *shot2* mutation leading to aberrant expression of downstream genes, which are defined as “read-in” genes in a previous study (45). These downstream read-in genes are expressed at a much lower level in heated *hot1-4* or in plants under normal growth conditions. The third and smallest cluster with seven genes contains genes down-regulated by heat in both *hot1-4* and *hot1-4 shot2*, most of which were not significantly different between the two genotypes under heat treatment conditions.

Next, we looked more closely at the effect of the *shot2* mutation on heat-regulated genes. The 5326 differentially heat-regulated genes (FDR < 0.01, |log2FC| >1) in *hot1-4* were traced for their differential expression (FDR < 0.01) in *hot1-4 shot2* upon heat treatment. We also looked at the differential expression (FDR < 0.01) of the same genes in *hot1-4 shot2* compared to *hot1-4* under heat treatment conditions (Fig. 3d). The heat map in Fig. 3d and pie charts in Fig. 3e (log2FC values are in Additional file 2: Table S7) show that most heat-regulated genes in *hot1-4* were differentially regulated in the same direction in *hot1-4 shot2* (2082 out of 2785 increased, 1934 out of 2541 decreased). Most notably, 422 out of 2785 heat-inducible genes were still heat-induced, but transcripts accumulated to lower levels in *hot1-4 shot2* (shown with an asterisk in the left panel of Fig. 3e), suggesting that CstF77 function is required for full induction of these genes by heat treatment or that the aberrant, incorrectly terminated transcripts are significantly more unstable than correctly processed transcripts. This group of genes includes many heat shock transcription factors (HSFs) and HSPs important for heat stress tolerance. Most genes in this group also show defects in transcription termination, resulting in reads downstream of the TTS similar to the *HSP17.8* gene (*HSP101* is shown as an example in Additional file 1: Fig. S6).

In summary, the RNA-Seq data demonstrate that *hot1-4 shot2* plants have reduced transcript levels of HSPs and HSFs compared to *hot1-4*. The fact that *hot1-4 shot2* is more tolerant to a 38 °C heat treatment than *hot1-4*, even with reduced expression of HSPs, suggests that reducing the toxicity of *hot1-4* is more important than expressing high levels of other HSPs and HSFs under these conditions.

### CstF77 is critical for a robust heat stress response

Our differential gene expression analysis suggests that the *shot2* mutation in CstF77 does not cause readthrough events in all transcripts. To investigate the genes whose correct transcription termination is dependent on CstF77, we developed a pipeline using established bioinformatic tools to systematically identify genes with transcriptional readthrough as illustrated in Fig. 4a. Based on the Araport11 annotation of the *A. thaliana* genome (46), TTS of every gene in the genome was assigned according to the Araport 11 annotation. Then, we used bedtools to intersect alignments for each of our 12 samples with the regions 100 bp upstream and downstream of the TTS of every annotated gene in the genome (47). Minimum overlap of a read to be counted for either region was set at 30%. Genes with mean read counts of less than 10 for the upstream regions in heat-treated *hot1-4* were removed because they were not likely to have readthrough reads due to low expression levels. A total of 589 genes passed this filter (Additional file 2: Table S8a). The ratios (R) of downstream to upstream read counts were calculated for all samples. Then, mean ratios (R) of the three biological replicates were calculated for each of the four experimental groups. Ideally, if a gene has a mean ratio close to one, it is more likely to have readthrough transcripts. The likelihood of readthrough transcription (LoR) for each gene in *hot1-4 shot2* compared to *hot1-4* under heat treatment was then calculated by subtracting the mean ratios of *hot1-4* from those of *hot1-4 shot2*. Again, if a gene has a LoR value close to one, it would be a strong candidate with readthrough transcription caused by the *shot2* mutation under heat treatment, although the real values can be even higher than one, due to fluctuation of sequencing depth along transcripts.

**Fig. 4.**
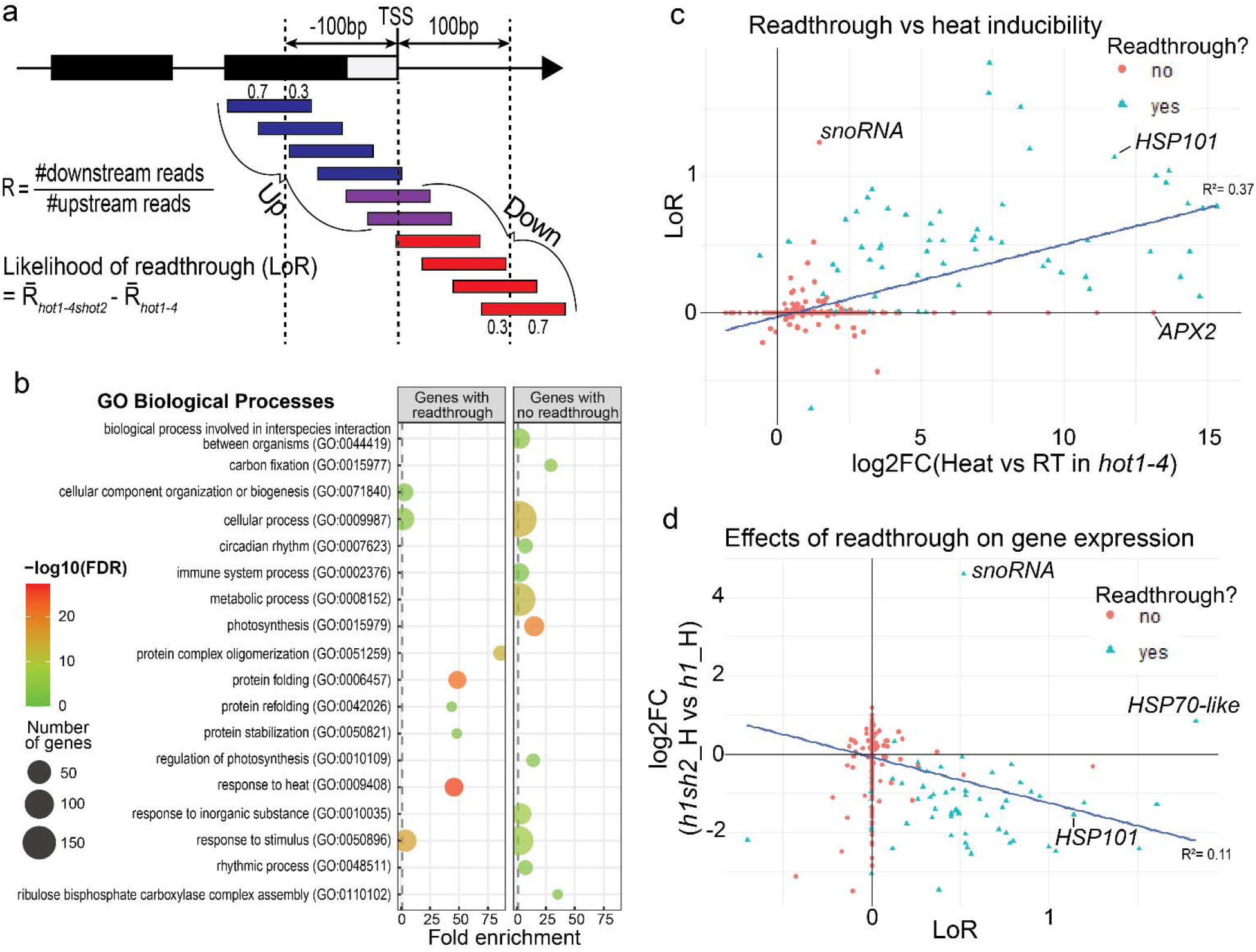
Highly heat-inducible genes are affected most by *shot2*. (a) Illustration of how the likelihood of readthrough transcription (LoR) is calculated. The number of reads that align 100 bp upstream and downstream of a transcription stop site (TTS) of a gene are counted. The ratios (R) of downstream read counts to upstream read counts are calculated and averaged for the three biological replicates of the four groups. LoR by *shot2* under heat stress is calculated by subtracting the mean ratios of *hot1-4* from the mean ratios of *hot1-4 shot2*. A putative gene model is depicted with exons in black rectangles, 3’-UTR in white rectangle and introns with a line. The blue and red reads overlap at least 30% with the upstream and downstream region, respectively, while the purple reads overlap with both regions. (b) Over-represented GO biological processes in genes with or without readthrough reads. The most representative and significant biological processes are shown with fold enrichment. The dot size indicates the number of genes associated with the process and the dot color gradient indicates the significance of the enrichment (−log10 (FDR-corrected P-values)). The vertical grey dashed lines represent a fold enrichment of 1.0. (c) 527 genes with at least 10 reads in 100bp upstream region of TTS in heated *hot1-4* samples were plotted to look at the relationship between heat inducibility (log2FC (H vs RT in *hot1-4*)) and LoR. A regression line with R^2^ value is overlayed on the graph to show the trend. H: Heat, RT: room temperature. (d) The same genes were plotted to look at the relationship between readthrough (LoR) and gene expression levels in *hot1-4 shot2* compared to *hot1-4* under heat treatment. *h1sh2_H: hot1-4 shot2* heated samples, *h1*_H: *hot1-4* heated samples.

All 589 genes that passed the filter of at least 10 reads in 100bp upstream region from TTS in heated *hot1-4* samples were also checked to determine whether they had real readthrough reads by directly examining BAM files with the Integrative Genomics Viewer (IGV) (GroundTruth column in Additional file 2: Table S8a). Seventy-six genes of the 589 genes had clear readthrough reads (55 with an LoR >0.1), while 505 genes did not (only 25 of them had an LoR >0.1). The remaining eight genes included rRNAs and noncoding RNAs with too many reads to be displayed in IGV. Gene ontology (GO) enrichment analysis of the 76 genes with readthrough reads revealed that GO terms such as response to heat and protein folding are highly enriched (Fig. 4b). On the other hand, GO enrichment analysis of the 505 genes with no readthrough reads showed enrichment of genes related to photosynthesis (Fig. 4b). Most genes in the latter category are highly expressed irrespective of heat treatment. During our analysis we also found that there were genes with mis-annotated TTS (34 out of 589). As an example, the TTS of the *HSFA2* gene is annotated more than 100 bp downstream from the actual TTS evident from the sequencing data. Thus, it has less than 10 upstream read counts and was filtered out of our analysis even though it has clear readthrough reads (Additional file 1: Fig. S7a). Therefore, the number of readthrough genes we identified is an underestimation.

To examine in detail the relationship of readthrough events and differential expression by heat treatment or by the *shot2* mutation under heat treatment, we included log2FC and FDR for each of the comparisons in the dataset. From 589 genes with at least 10 reads in 100bp upstream region of the TTS in heated *hot1-4* samples, we removed 62 genes that have a misannotated TTS or too many reads to be displayed in IGV or that reside on chromosome 2 where the alignment of reads is questionable due to mitochondrial genome duplication (Additional file 2: Table S8b). As shown in Fig. 4c, there is a clear positive relationship between heat inducibility and readthrough events, confirming that the *shot2* mutation mostly affects highly heat-induced genes. There were some exceptions to this trend including the ASCORBATE PEROXIDASE 2 (APX2) gene, which has a huge fold-increase (13.13 log2FC), but no readthrough reads. However, APX2 has a high fold increase with heat treatment because it is not expressed at all under normal growth conditions (Additional file 1: Fig. S7b); the absolute expression level of APX2 under heat stress is considerably lower compared to the expression levels of typical HSF or HSP genes.

Under heat treatment conditions, most genes with readthrough transcripts in *hot1-4 shot2* had lower transcript levels compared to the same genes in *hot1-4* (Fig. 4d), indicating that failure of polyadenylation and transcription termination negatively affects transcript abundance. One of the exceptions was a small nucleolar RNA (snoRNA, AT1G07897) which had a 4.6 log2-fold increase. This gene has readthrough reads and higher transcripts in *hot1-4 shot2* because it is located downstream of the *HSP17.6C* gene (Additional file 1: Fig. S7c). Many snoRNAs are present in our analysis likely due to their high expression levels. However, none of the other snoRNAs has readthrough transcripts, suggesting that maturation of snoRNAs is not dependent on CstF77.

It has been recently reported that heat stress can induce widespread transcriptional readthrough events (1,871 genes) in mouse fibroblast cells (48). To examine whether heat stress can have similar effects in *A. thaliana*, we applied our bioinformatics pipeline for identifying readthroughs to *hot1-4* samples with or without heat stress. Only 13 genes out of 589 genes showed readthrough reads, but at much reduced levels compared to readthroughs seen in *shot2* (Additional file 2: Table S9). The expression level of all the readthrough genes were increased by heat treatment and 12 of them also showed readthrough reads in heated *hot1-4 shot2* samples, suggesting that heat stress has limited effects on transcription termination in *A. thaliana*.

In summary, our bioinformatic analyses suggest that the function of CstF77 is critical for proper termination and full accumulation of highly heat-inducible transcripts, which is essential for a robust heat stress response.

### The *shot2* single mutant is more sensitive to heat stress

Defective transcription termination and polyadenylation are expected to negatively affect downstream processes including transcript stability, nuclear export, and translation, which could lead to more severe down-regulation at the protein level than at the transcript level. Therefore, it is important to examine the effects of *cstf77* mutations on HSP101 more quantitatively at the protein level. We performed immunoblot analysis to measure HSP101 reduction in *shot2* and plants with the partial loss-of-function *cstf77-1* allele. In the suppressed *hot1-4 shot2* plants, the mutant HSP101 (hot1-4) protein is reduced to 16.4% of the protein level in *hot1-4* (Fig. 5a), which is a larger decrease than the 34% decrease at the transcript level observed in the RNA-Seq analysis. This suggests that readthrough transcripts likely have further post-transcriptional defects in, for example, nuclear export and/or translation efficiency. In plants carrying the other mutant CstF77 allele, *cstf77-1*, the wild-type HSP101 protein level was also significantly reduced to 47% of the corresponding wild type, JU223 (Fig. 5a). This result suggests that the *cstf77-1* mutation causes a milder defect in the function of CstF77, and presumably milder heat stress phenotypes.

**Fig. 5.**
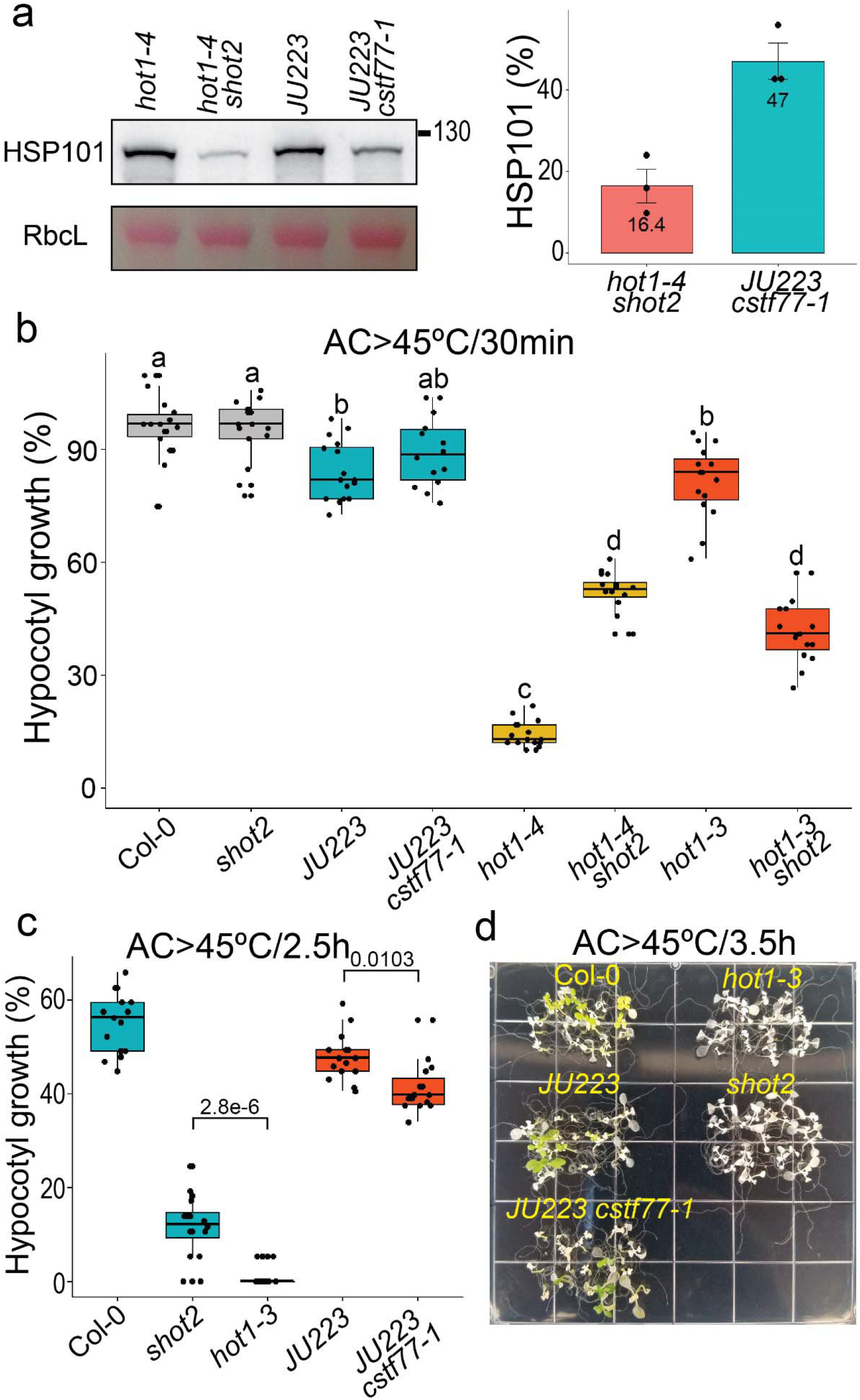
The *shot2* single mutant is more sensitive to heat stress. (a) HSP101 protein accumulation is reduced by mutations in the CstF77 gene. Total proteins were extracted after 38 °C for 3h followed by 1h recovery at a room temperature. A representative immunoblot is shown on the left. Ponceau S stained RbcL (Rubisco large subunit) is a loading control. Quantitative measurements of HSP101 protein level relative to each wild type from three independent biological samples are shown in a bar graph on the right. Error bars represent a standard deviation. (b) Heat stress tolerance was measured by hypocotyl elongation after heat stress treatment at 45 °C for 30 min following acclimation treatment (AC>45°C/30min). Different letters indicate significant differences (p<0.01) by one-way ANOVA followed by Tukey’s post hoc test. (c) Heat stress tolerance was measured by hypocotyl elongation after 45 °C for 2.5h following acclimation treatment (AC>45°C/2.5h). P-values from two-tailed t-tests of each comparison are shown in the graph. (d) Heat stress tolerance of light-grown seedlings was tested after 45 °C for 3.5h following acclimation treatment (AC>45°C/3.5h). The heat stress assays in (b), (c) and (d) were repeated more than once with similar results.

Since mutations in CstF77 cause reduction in many heat-inducible genes including *HSP101*, they are expected to impact heat stress phenotypes in other genetic backgrounds. To investigate the effects of *shot2* mutation on thermotolerance in the HSP101 null (*hot1-3*) or wild-type backgrounds, we replaced the *hot1-4* mutation by crossing the original *hot1-4 shot2* mutant with *hot1-3* or wild-type Col-0. After a relatively short (30 min) incubation at 45 °C following acclimation treatment (38 °C/90min, 22 °C/2h), the *shot2* single mutant does not exhibit any heat stress phenotype in a hypocotyl elongation assay (Fig. 5b). However, in the *hot1-3* background, the *shot2* mutation made plants more heat-sensitive to the 45 °C treatment (Fig. 5b). This result indicates that proper 3’ end maturation of transcripts other than *HSP101* is also required for thermotolerance. With more severe heat treatments (2.5 h or longer at 45 °C) (Fig. 5c and 5d), the *shot2* single mutant exhibited a strong heat-sensitive phenotype almost as severe as *hot1-3* plants in both hypocotyl elongation assays and light-grown seedling assays. The slightly better heat tolerance of *shot2* over *hot1-3* (Fig. 5d) may reflect residual HSP101 activity in the *shot2* mutant. Considering that the *shot2* mutant is affected not only in HSP101, but also in many other heat inducible proteins important for thermotolerance, these data highlight the dominant role of the HSP101 protein in the protection of plants from severe heat stress. Interestingly, the *cstf77-1* mutant plants with a HSP101 protein level at 47% of wild-type showed only slight heat sensitivity after 45 °C/2.5h treatment following acclimation (Fig. 5c). This result supports the conclusion that the relationship between heat resistance and HSP101 level is non-linear allowing a broad upper range of HSP101 protein level for effective tolerance to severe stress.

Taken together with the RNA-Seq results, our data demonstrate that proper 3’ mRNA maturation is necessary for the maximum expression of heat-inducible proteins and a robust heat shock response in plants.

## Discussion

We isolated and characterized a temperature sensitive mutation in CstF77 that reveals the importance of mRNA 3’ end formation in gene expression under heat stress conditions. We demonstrated that the loss of CstF77 activity in the *shot2* mutant under high temperature leads to defective transcription termination as evidenced by massive readthrough events mainly of highly heat-inducible genes, suggesting that the cleavage of mRNA 3’ ends is critical for a proper transcription termination. Furthermore, heat stress phenotypes of *shot2* mutants in *hot1-4, hot1-3* and wild-type backgrounds also reveal the dominant role of HSP101 in tolerance to severe heat stress.

CstF77 is an evolutionarily conserved protein from yeast, fruit flies, mammals and plants encoded in a single copy gene (6,9). The homologs in yeast (Rna14) and in fruit flies (suppressor of forked) are essential for viability (49,50). It is interesting that CstF77 in *A. thaliana* is reported to be non-essential for viability, but critical for normal auxin signaling and plant development (43). The T-DNA knockout mutant, *cstf77-2* showed shifts in polyadenylation sites from proximal to distal sites in most genes and differential expression of many stress-related genes under a normal growth condition (43). To our surprise, however, massive transcriptional readthrough events observed with our temperature-sensitive mutant were not reported in a transcriptomics study of the *cstf77-2* mutant under normal growth conditions. There could be several explanations for the lack of apparent transcription termination failure in *cstf77-2*, such as differences in RNA-Seq library prep (poly(A) selection versus ribosomal depletion used in our study), differences in the mutant alleles investigated (*cstf77-2* vs *shot2*) and different temperature conditions. Resolving this issue in future investigations would be important as it relates to basic understanding of the role of Cstf77 in mRNA 3’ end maturation of plants under normal and heat stress conditions.

Recently, cellular stresses including osmotic stress, heat stress, hypoxia, influenza A virus (IAV) infection, herpes simplex virus 1 (HSV-1) infection, and cancer have been demonstrated to disrupt normal transcription termination, leading to widespread readthrough transcription in yeasts and animals (45,48,51–55). IAV produces the viral non-structural protein1 (NS1), which binds to CPSF30 and presumably inhibits cleavage of pre-mRNAs, leading to readthrough transcription of host genes (56). However, molecular mechanisms of readthrough transcription in most stress conditions are not known. It is worth noting that heat shock induces readthrough transcription in 1871 genes in mouse fibroblast cells (48), but only in a limited number of genes (13 genes) in *A. thaliana*, highlighting differences in the effects of heat shock on transcription termination in the two organisms. The difference might lie on the milder heat stress treatment (38 °C for 3h) in our study with *A. thaliana*. More studies will be necessary to decide whether the effects of stresses on transcription termination are conserved in plants or specific to animal systems.

The effects of the temperature-sensitive *shot2* mutation on transcription termination tend to be stronger in highly heat-induced genes. This result is somewhat expected when we assume that the mutant protein has some residual activity in the stress condition used in our experiments. The residual activity of the shot2 mutant protein may be sufficient for 3’ end cleavage of low- to medium-expressed genes, but not enough for 3’ end maturation of highly expressed genes. However, it is difficult to explain why there is almost no effect on transcription termination of constitutively expressed genes such as photosynthesis-related genes, for example *LHCA4* (AT3G47470) (Additional file 1: Fig. S8). One explanation could be that transcription of these genes might be shut off at high temperature and the transcripts are so stable that they persist even after heat stress. However, some of these genes are significantly upregulated by heat stress (e.g. 1.92 log2FC for *LHCA4*), although at a much lower log2FC than HSP genes, indicating that genes such as *LHCA4* are being newly transcribed at high temperature, in which case some readthrough events would be expected. Further studies on dynamics of de novo mRNA synthesis and decay using innovative technologies such as 4-thiouridine labeling and nascent transcript sequencing (57) may provide insights on this phenomenon.

Most genes with readthrough transcription exhibit reduced transcripts in *hot1-4 shot2* compared to *hot1-4*, which likely results from increased mRNA decay due to defects in the poly(A) tail. Interestingly, HSP101 is reduced even more at the protein level (16.4% of WT) than at the transcript level (34% of WT), indicating that the mis-terminated RNAs may not be effectively translated. Consistent with this observation, previous studies with human and mouse cells show that readthrough transcripts remain in the nucleus and are not translated (45,48,55). Readthrough transcription in *shot2* also result in increased transcripts of genes downstream of genes where transcription is not terminated normally. We observed that these read-in genes have defects in splicing similar to results from a recent study in yeast where they depleted Nab2, which is required for proper 3’ end processing (58). Therefore, it is unlikely that transcripts of the read-in genes are transported and translated into proteins.

To identify putative readthrough genes, we built a custom bioinformatics pipeline that relies on robust and well-maintained software packages. This pipeline allowed us to whittle our list of several thousand differentially expressed genes down to those that were most likely to exhibit transcriptional readthrough. Our protocol can be readily modified for different systems and is freely available on GitHub (https://github.com/JDSwenson/Transcriptional-readthrough). When applied to our data, we successfully identified 55 readthrough genes with a LoR value >0.1 caused by the *shot2* mutation under the heat stress condition. This is no doubt an underestimation of the actual number of genes with readthrough. The biggest hurdle in finding a more accurate number originates from mis-annotation of TTS in the current *A. thaliana* genome annotation (Araport 11). Increasing the 100 bp window size of upstream and downstream region for calculation of LoR may allow detection of additional genes with mis-annotated TTS, but it could also increase the number of false-positives because reads belonging to a downstream gene could be counted as readthrough reads. Therefore, mis-annotated TTS in the current Araport 11 version need to be updated based on abundant RNA-Seq data available in various repositories for more accurate detection of readthrough genes.

The mechanism by which *shot2* suppresses the heat-sensitive *hot1-4* mutation is evident. Because the hot1-4 protein is toxic to plants, any mutation that reduces the activity of hot1-4 would suppress the heat-sensitive phenotype. In fact, many intragenic suppressors were reported previously where a second mutation in the *hot1-4* gene inactivates HSP101 function (29). The heat stress phenotypes of *shot2/cstf77* mutants in the HSP101 null mutant *hot1-3* and wild-type backgrounds strongly suggest the dominant role of HSP101 in the protection of plants from heat stress (Fig. 5). Heat stress in plants triggers complex transcriptional regulatory networks composed of many transcription factors (59,60). The *shot2* mutation causes defects in transcription termination resulting in transcriptional readthrough events and reduced expression of many heat-inducible transcription factors including *HSFA2* (−1.25 log2FC), *HSFB1* (−1.27 log2FC), *MULTIPROTEIN BRIDGING FACTOR 1C (MBF1C*, −1.82 log2FC and, *DEHYDRATION-RESPONSIVE ELEMENT BINDING PROTEIN 2A (DREB2A*, readthrough but not a significant fold-change). As a result, the heat shock response is severely dampened in *shot2* mutants. The fact that *hot1-4 shot2* is more thermotolerant than *hot1-4* emphasizes the toxicity of the hot1-4 protein, the reduced expression of which allows plants to overcome any negative effects of reduced expression in other HSPs. Once the toxic hot1-4 protein is removed, the *shot2* mutation makes both the HSP101 null *hot1-3* and wild-type plants more sensitive to heat stress. Surprisingly, the *shot2* single mutant, which accumulates around 16% of wild-type levels of HSP101 and less of most other HSPs, is more heat-tolerant than the *hot1-3* single mutant, which has no HSP101 but full expression of all the other HSPs. This result highlights the advantage of having at least some HSP101 over having full expression of all the other types of HSFs and HSPs for acquired thermotolerance.

## Conclusions

In summary, the identification and characterization of a temperature sensitive mutant in the mRNA 3’ end maturation factor CstF77 allowed us to elucidate the importance of mRNA 3’ end formation in a robust heat shock response of plants and the critical importance of HSP101 in the protein quality control network.

## Methods

### Plant materials, growth conditions and transformation

All *A. thaliana* mutants are in the Columbia-0 (Col-0) background, unless otherwise indicated: *hot1-3* (17), *hot1-4* (29). *cstf77-1* is in JU223 background (Landsberg *erecta* (L*er*) carrying an active FRIGIDA allele) (34,61). For mapping the *shot2* mutation, we introgressed *hot1-4* into L*er* background by backcrossing nine times.

Seeds were surface sterilized and plated on half strength Murashige and Skoog (MS) medium (62) supplemented with 0.5% sucrose and 0.8% phytoagar. Seeds were stratified at 4°C for 2 to 3 d. Plants were grown in controlled growth chambers (100 μmol m^−2^ s^−1^) on a 22°C/18°C, 16-h-day/8-h-night cycle.

For thermotolerance assays, 2.5-d-old dark-grown and 10- to 12-d-old light-grown seedlings were treated as described previously (17,63).

### Identification of the *shot2* mutation

Approximately 7500 homozygous seeds of *hot1-4* were mutagenized with ethyl methanesulfonate and 170,000 M2 seeds were screened for suppressor mutants as described previously (29). Briefly, M2 seeds were surface-sterilized and plated on agar medium containing 0.5% sucrose. 2.5-d-old dark-grown seedlings were treated at 38 °C for 2 h, and then the plates were returned to the dark at 22 °C for 2.5 d. Seedlings that showed increased hypocotyl elongation compared with *hot1-4* were rescued by growth under light for approximately 1 week before being transplanted to soil. M3 plants were retested for tolerance to 38 °C for 2-3 h to confirm the suppressor phenotype.

Sequencing of the corresponding genomic region of *HSP101* gene confirmed that the mutants still carried the original *hot1-4* mutation and no other mutations in the *HSP101* gene.

To identify the causal mutation of the *shot2* suppressor phenotype, a mapping population was created by crossing *hot1-4 shot2* with the L*er* ecotype harboring the *hot1-4* mutation that had been created by introgression. 100 individual F2 plants with the suppressor phenotype were selected for seeds. The F3 plants were grown and pooled for next-generation Illumina sequencing in the Genome Access Technology Center at the McDonnell Genome Institute in St. Louis, Missouri.

The genomic *CstF77* DNA construct for complementation of the *shot2* mutation was generated by amplifying the appropriate DNA fragment (9009 bp) including the promoter (2032 bp upstream of the start codon) and terminator (913 bp downstream of the stop codon) regions with primers CSTF-3 and CSTF-4 (primer sequences are in Additional file 2: Table S10), and cloning the fragment into the entry vector pCR8/GW/TOPO (Invitrogen). The fragment was then inserted into the pMDC123 binary vector carrying Basta resistance (Curtis and Grossniklaus, 2003) by Gateway LR cloning (Invitrogen). Transformation of the construct along with the pMDC123 empty vector as a control into *hot1-4 shot2* was performed by floral dipping (Clough and Bent, 1998). T1 Transgenic plants were selected on half strength MS media containing 0.5% sucrose, 0.8% phytoagar and 10mg/L BASTA in the dark and tested for complementation by hypocotyl elongation assays on fresh media without BASTA as described previously (17,63).

### SDS-PAGE and immunoblot analysis

Total protein extracts were prepared by grinding frozen plant tissues in a 1.5-mL microtube with SDS sample buffer (60 mM Tris-HCl, pH 8.0, 60 mM DTT, 2% SDS, 15% Sucrose, 5 mM e-amino-N-caproic acid, and 1 mM benzamidine). Protein concentration was measured with a Coomassie Brilliant Blue binding assay (64). Protein samples were separated by SDS-PAGE and blotted onto nitrocellulose membrane for immunoblot analysis. Protein blots were stained with Ponceau S reversible protein stain (Sigma) to check transfer efficiency and equal protein loading. After the stain was removed, blots were probed with antibodies against HSP101 at 1:5000 (17), cytosolic HSP70s at 1:5000 (AS08 371, Agrisera), cytosolic small HSPs (HSP17.6 Class I) at 1:1000 (65) and cytosolic glyceraldehyde 3-phosphate dehydrogenase (GAPC) at 1:10000 (66). Blots were incubated with anti-rabbit IgG secondary antibodies conjugated with horseradish peroxidase (GE Healthcare) and visualized by enhanced chemiluminescence (Thermo Scientific Pierce ECL Western Blotting Substrate) using a G:Box iChemi XT(Syngene).

### RT-PCR and RNA-Seq experiments and analysis

For RT-PCR, Col, *hot1-4* and *hot1-4 shot2* seedlings grown on agar plates until stage 1.02 (67) were treated with or without heat stress at 38 °C for 3h in darkness followed by 22 °C for 1h in light. Total RNA was extracted using Trizol (LifeTechnologies) according to the manufacturer’s instructions. 2 μg of total RNA was reverse transcribed with random primers using SuperScript II (Invitrogen) according to the manufacturer’s instructions. Primer sequences are given in Additional file 2: Table S10.

For RNA-Seq samples, the *hot1-4* and *hot1-4 shot2* seedlings grown on agar plates until stage 1.02 (67) were treated with or without heat stress at 38 °C for 3h in darkness followed by 22 °C for 1h in light. Total RNA was extracted using a RNeasy Plant Mini kit (Qiagen, 74904). All samples were treated with DNase I using a RNase-Free DNase Set (Qiagen, 79254) and quantified. RNA samples prepared from three independent biological replicates were used for subsequent analysis. 7 μg total RNA was used for rRNA depletion with RiboMinus (Invitrogen, A10838-08) according to the manufacturer’s instructions. 100 ng of rRNA-depleted RNA was subsequently used for library preparation using NEBNext Ultra Directional RNA Library Prep Kit for Illumina (New England Biolabs, E7420S) according to the manufacturer’s protocol. Multiplex sequencing was performed on a NextSeq 500 (Illumina), and 75-bp-long single-end reads were obtained. The reads were mapped to the reference Arabidopsis genome (The Arabidopsis Information Resource 10) using HISAT2 (44) and counted using the htseq-counts script in the HTSeq library (68). Differential gene expression analysis was performed using the R package EdgeR (69). Search for genes with readthrough events were performed as described in the results section and scripts are housed on GitHub (https://github.com/JDSwenson/Transcriptional-readthrough).

Gene Ontology (GO) enrichment analysis and visualization of enriched GO terms were performed as described in Bonnot et al. (70) using Panther (71), REVIGO (72) and ggplot2 R package (73).

## Declarations

### Ethics approval and consent to participate

Not applicable

### Consent for publication

Not applicable

### Availability of data and materials

All whole-genome sequencing data from this study have been deposited in the NCBI under the accession number PRJNA772009 (https://www.ncbi.nlm.nih.gov/bioproject/PRJNA772009). The RNA sequencing datasets have been deposited in NCBI Gene Expression Omnibus with the accession number GSE185225 (https://www.ncbi.nlm.nih.gov/geo/query/acc.cgi?acc=GSE185225).

### Authors’ contributions

MK, FM, and EV designed the experiments, MK and FM performed experiments, MK and JS designed and completed the bioinformatics analysis. MK wrote the manuscript with input from all other authors. All authors read and approved the final manuscript.

## Acknowledgements

We thank Caroline Dean (John Innes Center. U.K.) for *cstf77-1* seeds and acknowledge services of the Genomics Resource Laboratory RRID:SCR_017907 of the University of Massachusetts Amherst Institute of Applied Life Sciences Cores with support from the Massachusetts Life Sciences Center. Research was supported by grants from the National Science Foundation (IOS 1354960), the Department of Energy Biosciences program (DE-FG02-99ER20338) and the Massachusetts Life Sciences Center to EV. JS acknowledges support from the Center for Models to Medicine of the University of Massachusetts Institute of Applied Life Sciences. We thank Peter Chien and Courtney Babbit for guidance in designing the bioinformatics analysis and Madeline Bartlett for bioinformatics help with *shot2* mapping.

