## Supplemental Figures for "A temperature sensitive mutation in the CstF77 subunit of the polyadenylation complex reveals the critical function of mRNA 3’ end formation for a robust heat stress response in plants"

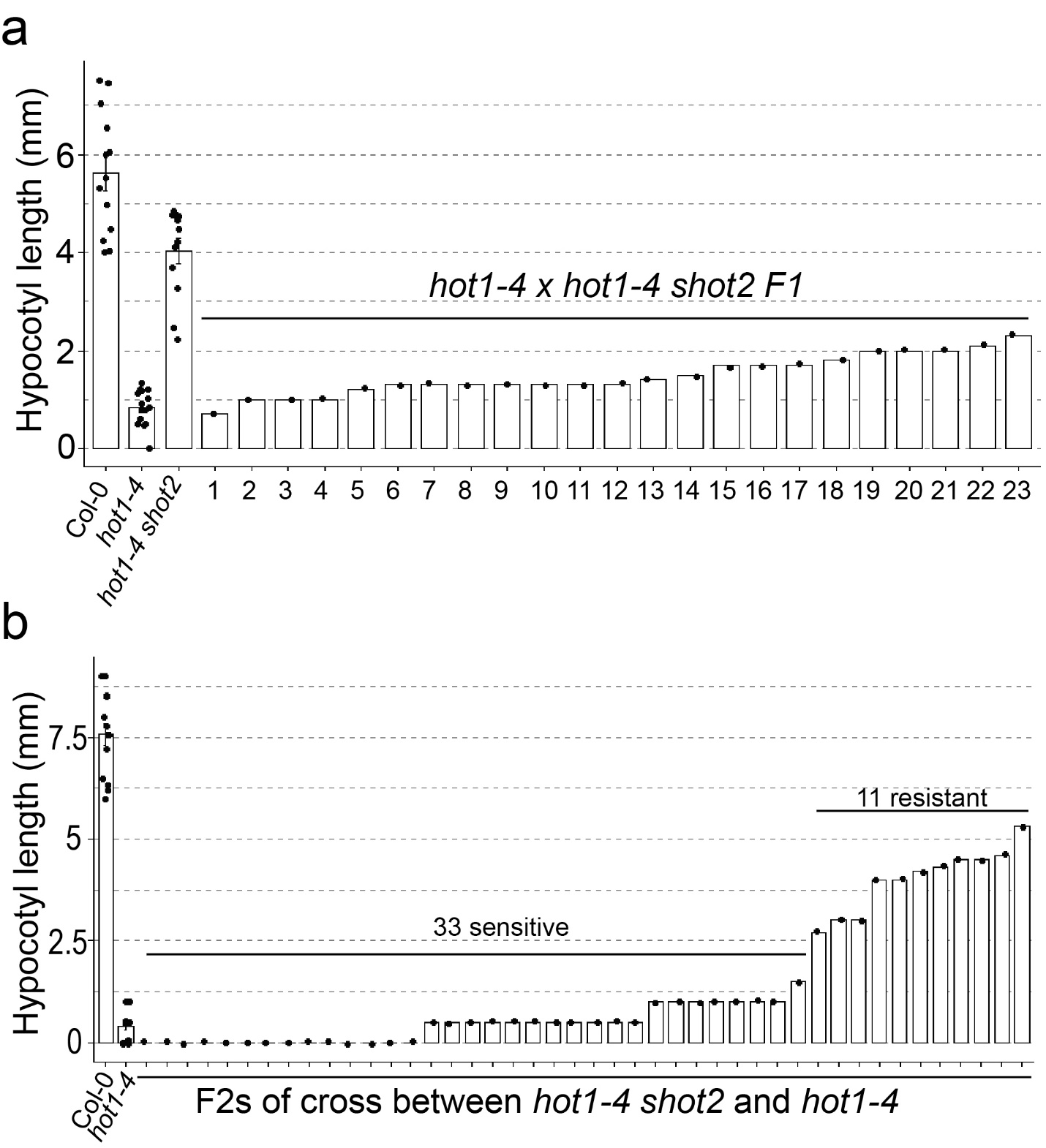

Fig. S1. The *shot2* mutation segregates as a single recessive locus. (a) 23 F1 and (b) 44 F2 seedlings of a cross between *hot1-4* and *hot1-4 shot2* were tested for heat tolerance by the hypocotyl elongation assay after heat treatment at 38 °C for 3h.

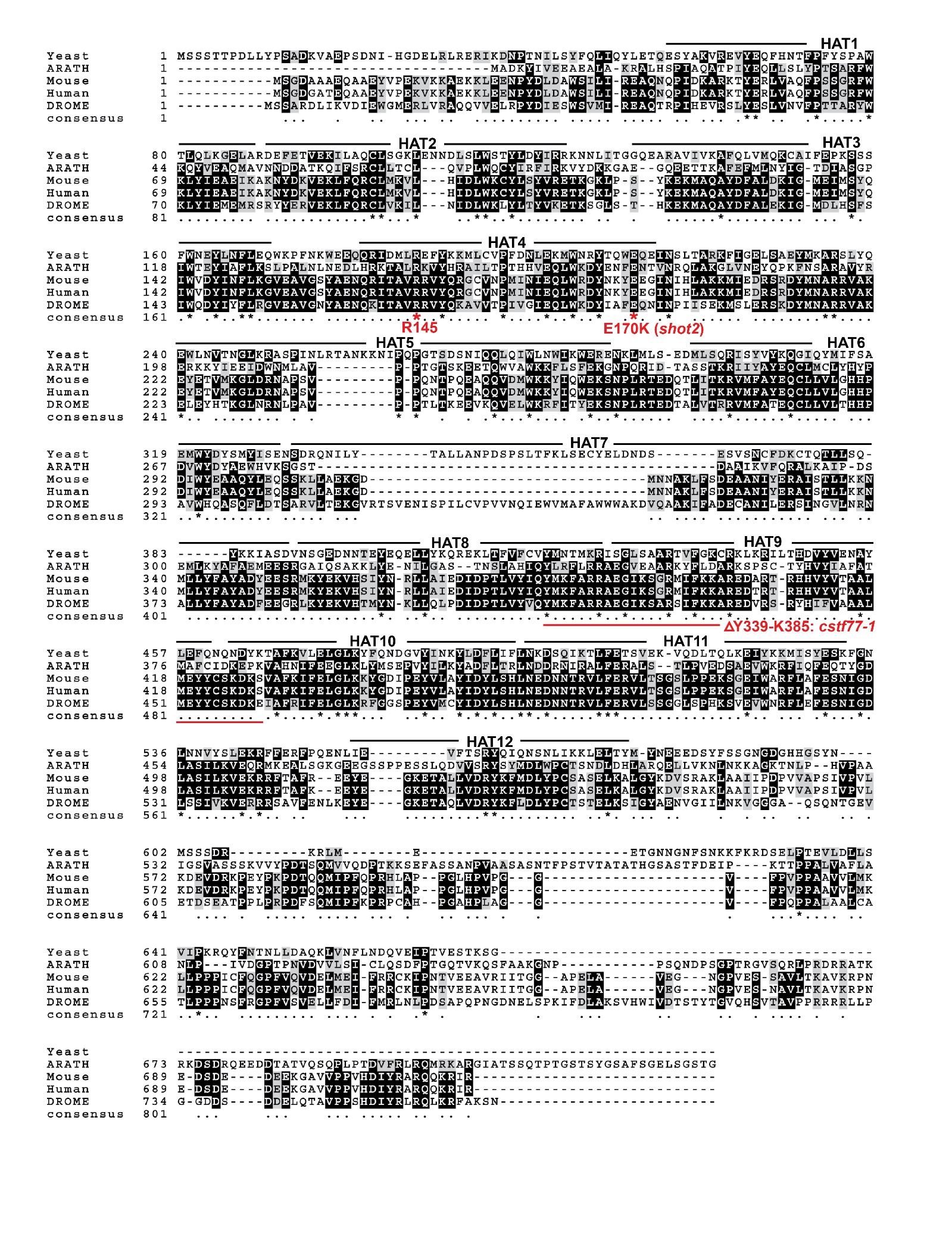

Fig. S2. Protein sequence alignment of CstF77 proteins from *Saccharomyces cerevisiae* (Yeast, Uniprot P25298), *Arabidopsis thaliana* (ARATH, Uniprot Q8GUP1), *Mus musculus* (Mouse, Uniprot Q99LI7), *Homo sapiens* (Human, Uniprot Q12996) and *Drosophila melanogaster* (DROME, Uniprot P25991). Multiple sequence alignment was performed using Clustal Omega (<https://www.ebi.ac.uk/Tools/msa/clustalo/>). The HAT motifs are shown as annotated in Uniprot database for *A. thaliana* CstF77. The *shot2* mutation (E170K), R145 (Hydrogen bond partner of E170) and *cstf77-1* mutation are shown in red. Identical residues are highlighted in black and similar residues in gray.

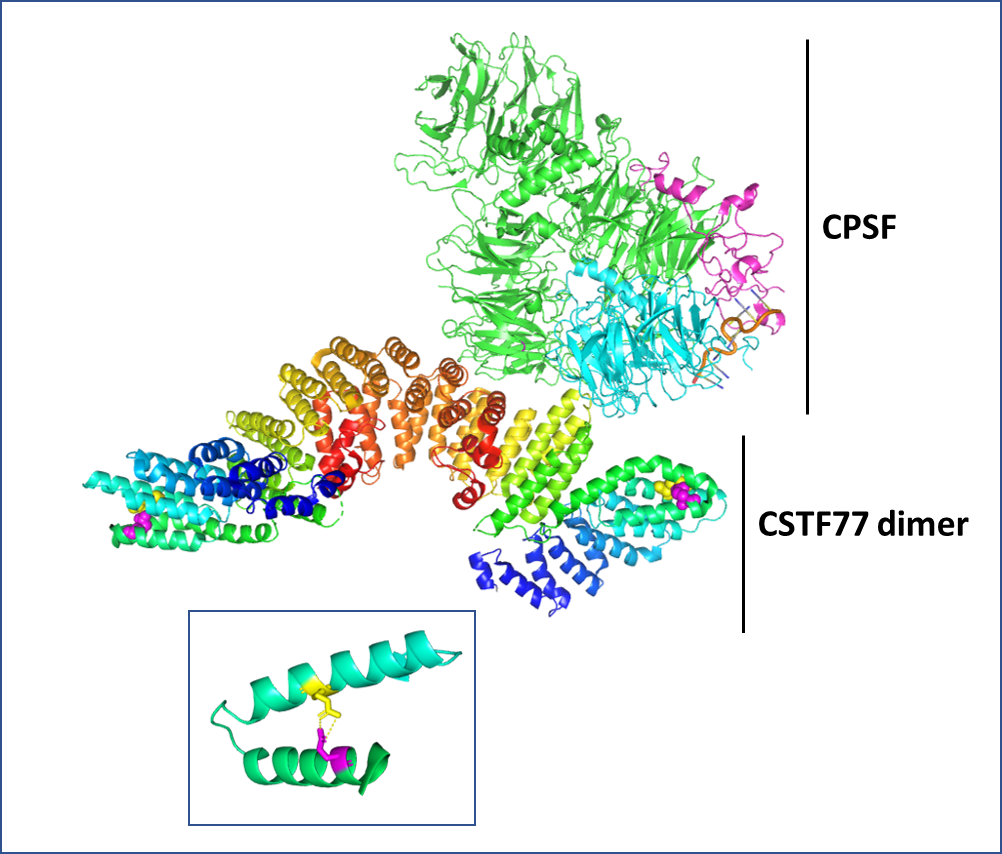

Fig. S3. Location of the *shot2* mutation on the structure of human Cstf77. Structure from PDB 6uro (Zhang et al 2020). CPSF subunits shown are CPSF160 (green), WDR33(cyan) and CPSF30 (pink) with RNA (AAUAAAACA, orange). Cstf77 dimer is shown in rainbow color from blue to red, N- to C-terminal. Note that the flexible C-terminal residues from aa 550-717 of human Cstf77 do not appear in the crystal structure due to their flexibility. Magenta spheres show position of residue mutated in *shot2*, E194, corresponding to E170 in *A. thaliana*. Yellow spheres show position of R169, corresponding to R145 in *A. thaliana*. Both residues are 100% conserved in Cstf77 sequences (Fig. S2). Inset: Illustrates H-bonding between E194 and R169.

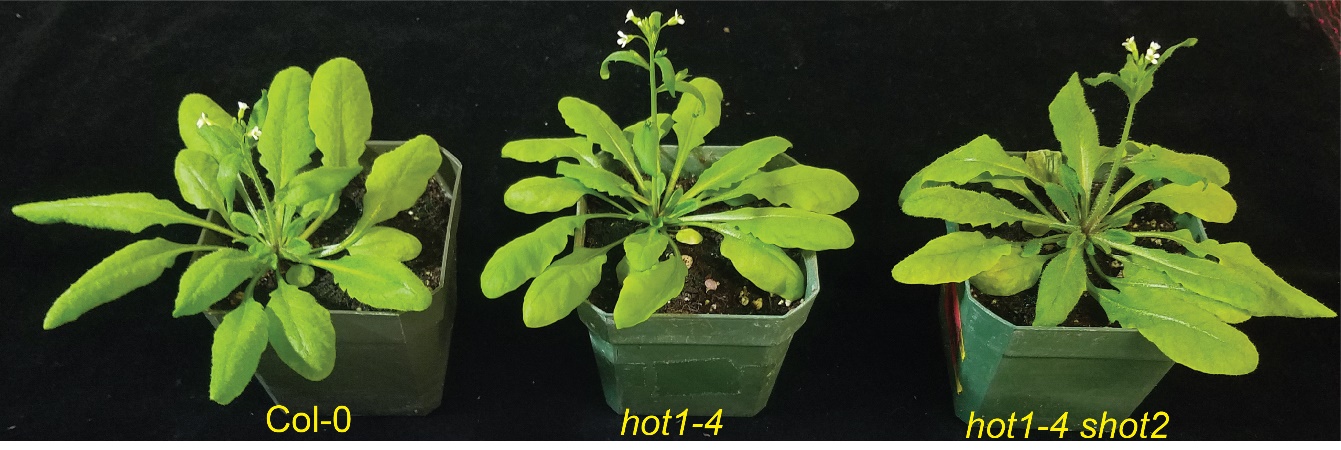

Fig. S4. *shot2* mutant plants grow like wild-type plants under a normal growth condition. Plants were grown under a long-day condition (16h light/8h dark) for 39 days.

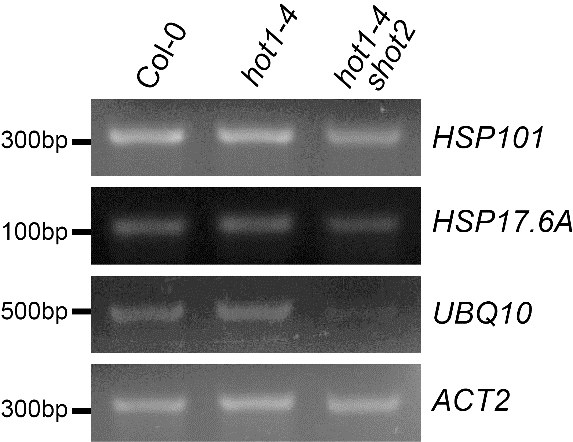

Fig. S5. Semi-quantitative RT-PCR analysis of the expression levels of *HSP101*, *HSP17.6A*, *UBIQUITIN10* (*UBQ10*), and *ACTIN2* (*ACT2*) in wild type Col-0, *hot1-4* and *hot1-4 shot2* seedlings subjected to heat stress at 38 °C for 1.5h followed by 22 °C for 1h. The experiments were repeated twice with similar results.

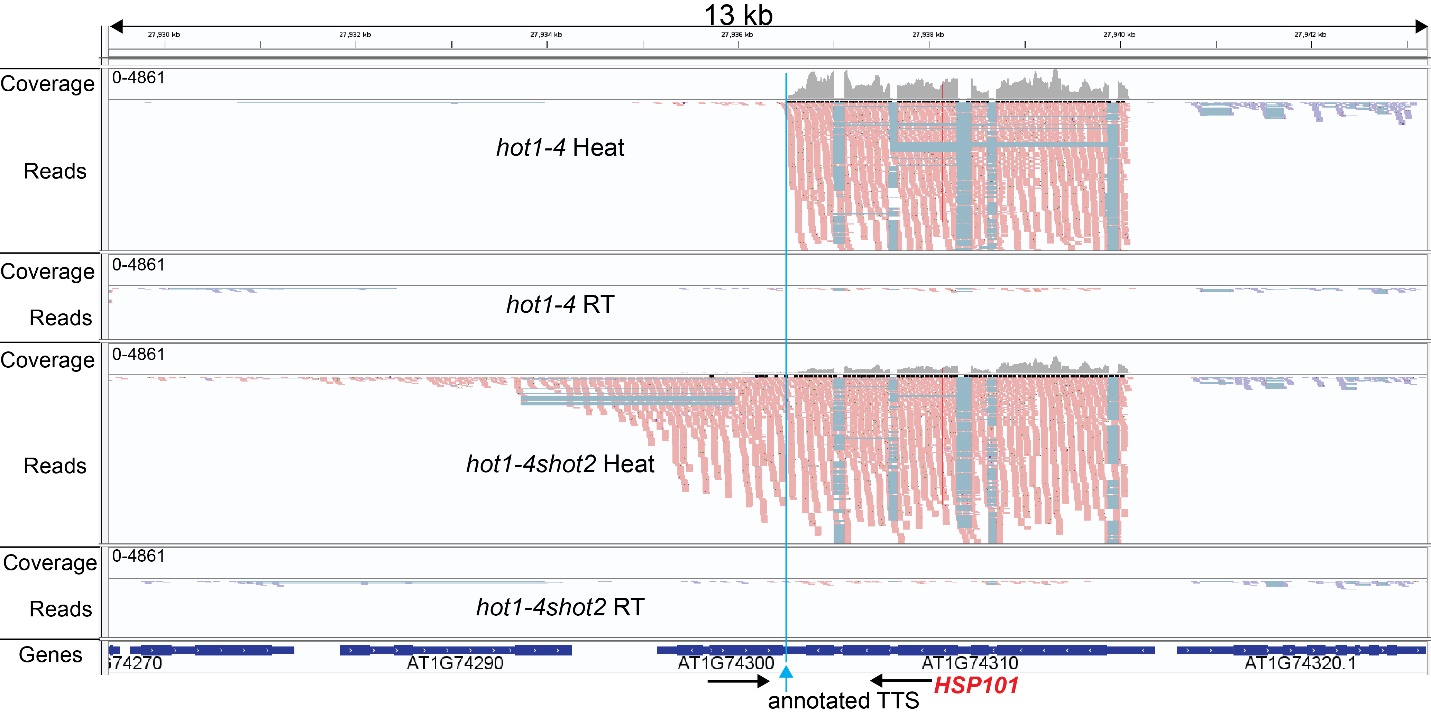

Fig. S6. An IGV (Integrative Genomics Viewer) snapshot of *HSP101* locus shows readthrough reads downstream of transcription termination site (TTS) in a heat-treated *hot1-4 shot2* sample. The orientation of genes is denoted by black arrows. The TTS of *HSP101* is indicated by a vertical blue line and a blue arrow. The red vertical lines indicate the *hot1-4* mutation.

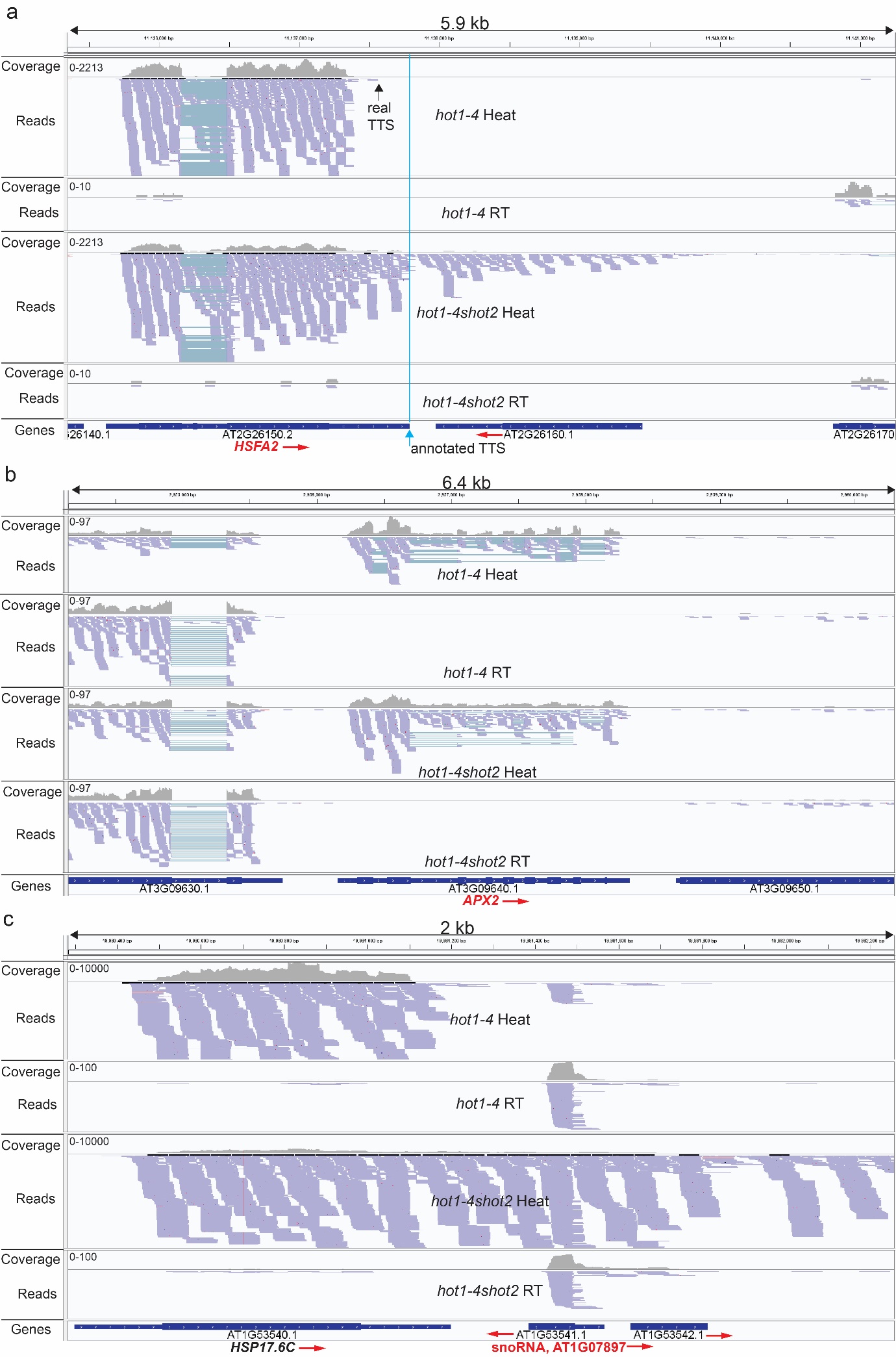

Fig. S7. An IGV snapshot of RNA-Seq reads mapped to *HSFA2, APX2 and AT1G07817 (snoRNA)* loci. The orientation of genes is denoted by red arrows. (a) *HSFA2* locus shows readthrough reads and a misannotated TTS in a heat-treated *hot1-4 shot2* sample. The actual TTS (indicated with a black arrow) is located more than 100 bp upstream from the annotated TTS (indicated with a blue arrow). (b) Highly heat inducible *APX2* does not have readthrough reads. Note that the read coverage range of *APX2* (0-97), which is significantly lower than other heat-inducible genes with readthrough reads (e.g. *HSFA2*, *HSP17.6C* and *HSP101*). (c) A snoRNA (AT1G07897) appears to have high amount of readthrough reads because of its location downstream of *HSP17.6C* with abundant readthrough reads. Note that the snoRNA overlaps with AT1G53541 which runs in the opposite direction, but with no expression.

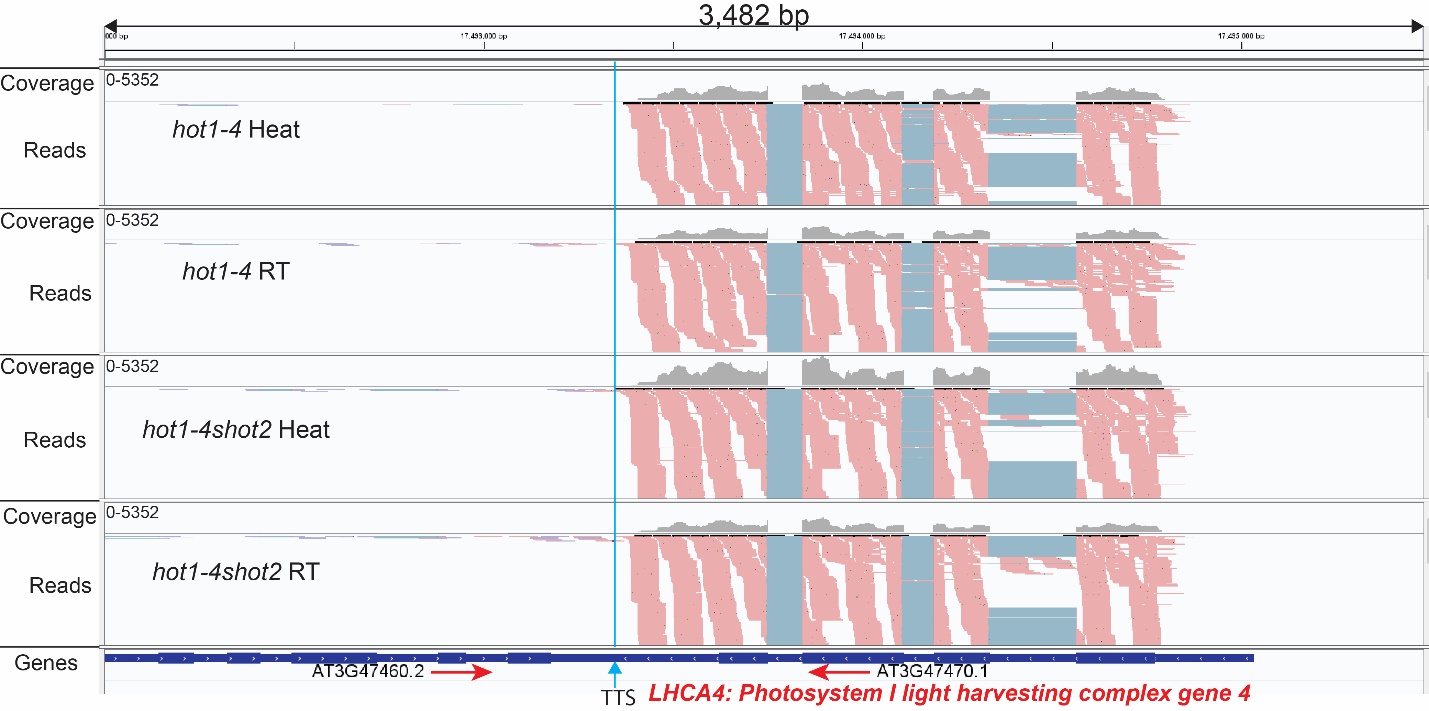

Fig. S8. An IGV snapshot of *LHCA4* locus shows no significant readthrough reads. The orientation of the gene is denoted by red arrows. The TTS of *LHCA4* is indicated with a blue line and a blue arrow.
