## Supplemental Table 1 for "A temperature sensitive mutation in the CstF77 subunit of the polyadenylation complex reveals the critical function of mRNA 3’ end formation for a robust heat stress response in plants"

Table S1. *SHOT2* candidate genes on chromosome 1. Frequency of alternative nucleotides at specific sites on chromosome 1 in the region of interest (Position) were identified from sequencing heat-tolerant F3 pools from crosses between *hot1-4* L*er* and *hot1-4 shot2* Col-0. Alt. Freq: Frequency of alternative nucleotides. Number of reads containing a changed nucleotide over the number of reads containing the reference nucleotide.

| Position | Reference | Change | Alt. Freq | Gene_ID | Gene_name | Bio_type | Effect | old_AA/new_AA | Old_codon/New_codon |
| --- | --- | --- | --- | --- | --- | --- | --- | --- | --- |
| 5525082 | C | T | 70/74 | AT1G16120 |  | downstream |  |  |  |
| 5630020 | C | T | 76/76 | AT1G16489 | AT1G16489 | ncRNA | EXON: exon_1_5629982_5630118 |  |  |
| 5630020 | C | T | 76/76 | AT1G16490 | ATMYB58 | protein_coding | NON_SYNONYMOUS_CODING | E/K | Gag/Aag |
| 5786547 | C | T | 74/77 | AT1G16916 |  | downstream |  |  |  |
| 5880320 | C | T | 68/68 | AT1G17210 |  | 3'utr |  |  |  |
| 6027929 | C | T | 70/70 | AT1G17530 | ATTIM23-1 | protein_coding | SYNONYMOUS_CODING | A/A | gcC/gcT |
| 6108302 | C | T | 85/85 | AT1G17750 | PEPR2 | protein_coding | SYNONYMOUS_CODING | S/S | tcC/tcT |
| 6115058 | C | T | 76/77 | AT1G17760 | CSTF77 | protein_coding | NON_SYNONYMOUS_CODING | E/K | Gaa/Aaa |
| 6137391 | C | T | 103/104 | AT1G17830 | DUF789 | protein_coding | STOP_GAINED | W/* | tgG/tgA |
| 6467736 | C | T | 68/68 | AT1G18750 |  | intron |  |  |  |
| 6835936 | C | T | 97/97 | AT1G19780 | ATCNGC8 | protein_coding | NON_SYNONYMOUS_CODING | R/K | aGa/aAa |
| 6912984 | C | T | 27/27 | AT1G19910 |  | upstream |  |  |  |
